# Hemispheric asymmetries in auditory cortex reflect discriminative responses to temporal details or summary statistics of stationary sounds

**DOI:** 10.1101/2023.08.03.551829

**Authors:** Martina Berto, Patrick Reisinger, Emiliano Ricciardi, Nathan Weisz, Davide Bottari

## Abstract

The processing of stationary sounds relies on both local features and compact representations. As local information is compressed into summary statistics, abstract representations emerge. Whether the brain is endowed with distinct neural architectures overseeing such computations is unknown.

In this magnetoencephalography (MEG) study, we employed a validated protocol to localize cortical correlates of local and summary representations, exposing participants to triplets of synthetic sound textures systematically varying for either local details or summary statistics. Sounds also varied for their sound duration, specifically short (40ms) or long (478ms). Results revealed clear distinct activation patterns for local features and summary statistics changes. Such activations diverged in magnitude, spatiotemporal distribution, and hemispheric lateralization. For short sounds, a change in local features, compared to summary statistics, predominantly activated the right hemisphere. Conversely, for long sounds, a change in summary statistics elicited higher activation than a change in local features in both hemispheres.

Specifically, while the right auditory cortex was responding more to changes in local features or summary statistics depending on sound duration (short or long, respectively), the left frontal lobe was selectively engaged in processing a change in summary statistics at a long sound duration. These findings provide insights into the neural mechanisms underlying the computation of local and summary acoustic information and highlight the involvement of distinct cortical pathways and hemispheric lateralization in auditory processing at different temporal resolutions.

**Significant Statement:** We revealed hemispheric specializations for auditory computations at high (local) and low (summary statistics) temporal resolutions. The right hemisphere was engaged for both computations, while the left hemisphere responded more to summary statistics changes. These findings highlight the multifaceted functions of the right hemisphere in capturing acoustic properties of stationary sounds and the left hemisphere’s involvement in processing abstract representations.

## Introduction

Naturalistic sounds are processed at low and high temporal resolutions, depending on the amount of information they encompass. When sounds are short, the processing of fine-grained temporal modulations (local features) is crucial for detecting transient changes in the entering sound waves. As the amount of information increases, retaining more and more features becomes challenging for the system, which must rely on compact representations. For specific categories of sounds, it is possible to mathematically describe such compact summary representations. For instance, stationary sounds (e.g., sound textures, such as fire, rain, and typewriting) are characterized by repetition over time of similar events. Thus, their summary representation can be described by a set of average statistics (envelope mean, variance, and skew; envelopes correlations, modulation power, and its cross-channel correlations; (McDermott & Simoncelli, 2011). Average summary statistics are spectrotemporal measurements computed over time from local features; they approximate the outcome of computations occurring in the periphery of the system and are encoded by both primary and post-primary auditory cortices (McDermott & Simoncelli, 2011; Giordano et al., 2023). The amount of acoustic information is crucial for the emergence of summary statistics from local features; thus, the engagement of one computational mode (i.e., local features processing) or the other (summary statistics) strictly depends on sound duration (McDermott et al., 2013).

The functional role of local features and summary statistics in the processing of stationary sounds has been evaluated in humans through computational synthesis approaches (Berto et al., 2021, 2022; McDermott & Simoncelli, 2011; McWalter & McDermott, 2019; Norman-Haignere & McDermott, 2018; Zuk et al., 2020); for instance, it was possible to disentangle the processing of local features from summary statistics by creating synthetic sounds with the same auditory statistics but different acoustic details (Berto et al., 2021, 2022; McDermott et al., 2013). Specifically, a previous study employed this approach to isolate neural signatures of local features and summary statistics of sound textures (Berto et al., 2022). In this study, brain activity was recorded using electroencephalography (EEG) while participants were exposed to streams of sounds comprising changes based on local features or summary statistics and the duration of the sound excerpt could be short, medium, or long to manipulate the reliability of local features or summary statistics processing. The results showed a clear dissociation: when sounds were short, a change in local features elicited higher evoked responses. In contrast, when sounds were long, a greater evoked response was measured for changes in summary statistics compared to local features. The two effects emerged over similar, but not identical, scalp locations, possibly indicating different sources. Moreover, different oscillatory profiles were associated with local or summary changes, with local features being encoded by faster oscillations than summary statistics (Berto et al., 2022). Interestingly, these results emerged without an explicit task. Thus, we could hypothesize that sound discrimination occurring at high or low temporal resolutions is automatically driven by selective stimulus properties. For instance, the discrimination of brief sound excerpts may rely on broadband impulse amplitude modulations, while the discrimination of long sounds may rely on spectral content averaged over time.

While it was possible to distinguish local features or summary statistics computations by neural activity, the limited spatial resolution of EEG did not allow localizing these effects on the cortical surface. Thus, it remained unknown whether the brain developed distinguished neural architectures devoted to the processing of fine-grained information compared to summary one.

Previous studies revealed an auditory pathway parcellation for temporal and spectral selectivity (i.e., Santoro et al., 2014). Hemispheric asymmetries were also observed (e.g., Flinker et al., 2019; Neophytou et al., 2022; Zatorre & Belin, 2001) with some studies suggesting left hemisphere sensitivity to temporal modulations and right hemisphere sensitivity to spectral content (Zatorre & Belin, 2001) and others suggesting greater left hemispheric selectivity for spectral modulations and weaker activations in the right hemisphere (Flinker et al., 2019). Contradictory findings likely stem from task and stimulus differences. Systematic synthesis approaches may overcome such limitations, and address specific computations hinged on attributes directly embedded in sounds (e.g., summary statistics). Moreover, they provide means to create sounds that are distinguishable at high, but not low, temporal resolution, and vice-versa (McDermott et al., 2013). This provides a compelling methodology for objectively disentangling the response to local features from summary statistics and testing whether auditory specializations exist in the context of local and summary processing.

In the current study, we leveraged the high spatial resolution of the magnetoencephalography (MEG) to provide reliable source estimates of the automatic processing of local and summary acoustic information, without losing temporal precision. We implemented a validated protocol (see Berto et al., 2022) and used short (40ms) and long (478ms) sounds presented in separate continuous streams at a constant rate (one sound every half a second). Within each stream, sounds appeared as a series of triplets, with the first two being identical and the third varying for its local features or summary statistics, depending on the experimental context. Importantly, sounds were extracted from a pool of different synthetic sound textures (Berto et al., 2022) and were never the same between participants and sound streams; however, while using different sounds excerpts, we always ensured that, within each experiment, repeated and novel sounds in the triplets systematically varied for either local features or summary statistics (see Material and Methods). This allowed us to specifically investigate the computation of interest while generalizing it to many different stationary sounds, disregarding their idiosyncratic properties (i.e., frequency, pitch, timbre, and amplitude of single sound excerpts).

Furthermore, this presentation scheme permits controlling for expectancy effects, as the novel sound always occurs as a third element in the triplet. Finally, creating regularities and then disrupting them (by presenting two identical sounds followed by a different one) permits the measurement of discriminative responses driven by local and summary computations without an explicit task.

## Material and Methods

### Participants

24 healthy individuals voluntarily participated in the investigations. Specifically, 8 females, 2 right-handed, with a mean age of 35.42 years (std=13.37, range=20-63). All participants reported no neurological impairments and normal hearing, which was confirmed by performing an audiogram before the experiment. Participants signed informed consent and were paid 10€ per hour for their contribution. The study was approved by the Ethics Committee of the Paris-Lodron-University of Salzburg, in accordance with the Declaration of Helsinki.

### Data exclusion

Two participants were excluded from the analysis, leading to a final sample size of 22 subjects, 7 females 7, 2 left-handed, with a mean age of 35.09 years (std=13.92, range=20-63). One participant was excluded due to technical problems during the recording; the other participant’s data was rejected because of co-registration failure (for more details on the latter, see the Source Reconstruction paragraph).

### Stimuli

Experimental sounds were a sub-selection of the ones used in the previous EEG study (Berto et al., 2022) and behaviorally validated by Berto et al. (2021) with the same protocol by McDermott, Schemitsch, and Simoncelli (2013). They represented short snippets (either 40 or 478ms long) extracted from 5s synthetic sound textures. Each synthetic sound was produced by using the computational texture model designed by McDermott and Simoncelli (2011; see Extended Methods in Supplementary Material). A set of spectrotemporal statistics was extracted from 54 original recordings of 7s natural sound textures (i.e., rain, applause, waterfall; see the complete list in Table S1, Supplementary Material) and imposed on four different 5s white noise samples (Figure S1; for more details see the Extended Methods in Supplementary Material). The outcomes were different synthetic sounds differing for their local temporal features (which depends on the input white noise) but including the same summary statistics (Figure S1). All the four synthetic exemplars were cut into excerpts of 40 or 478ms to which a 20ms half-hann window was applied (10ms at the beginning and 10ms at the end) to reduce edge artifacts. Excerpts were numbered according to their position along the 5 seconds signal (i.e., excerpt 1 occurs from 1 to 40ms, excerpt 2 from 41 to 80ms and so on). We will refer to this number as their positional index (see below). All excerpts were equalized to the same root mean squared amplitude (rms) of 0.1 and had a sampling rate of 20kHz.

### Data acquisition

The brain activity was recorded using a 306-channel whole head MEG system comprising 204 planar gradiometers and 102 magnetometers (Neuromag TRIUX, Elekta). Signal was recorded with a sampling rate of 1000Hz and filtered online at 0.1Hz. Prior to recording the MEG, digital head-shape points (HSP) were measured on the scalp using a Polhemus Fastrak system (Polhemus, Colchester, VT). Approximately 500 points for subject were acquired, including three fiducials (nasion and left and right preauricular points). Three electrodes were applied to measure ocular and cardiac artifacts: two EOGs (vertical and horizontal channels attached above and lateral to the right eye), and one ECG.

Participants were then accompanied in the magnetically shielded room (AK3B, Vacuumschmelze, Hanau, Germany) where the MEG system was located and sat comfortably in front of a screen located at a ∼110cm distance. The experimental instructions and a fixation cross were back projected on the translucent screen by a Propixx DLP projector (VPixx Technologies, Canada). Finally, participants were wearing MEG-compatible insert earphones.

### Experimental protocol

The experiment consisted of 4 blocks of 5.4min each. Participants listened to streams of sounds but were instructed to ignore them and simply press a button when a target one occurred. The target sound was very infrequent (maximum three sounds per block) and consisted of a 50ms pure tone with a frequency of 2200Hz, an amplitude of 50dB, and a sampling rate of 20kHz.

Within each block, 648 sounds were presented at a continuous stimulation rate (one sound every 500ms). The stream in each block contained sounds of one specific length, either short (40ms) or long (478ms), so that the duration of the stream was kept constant, but the amount of acoustic information changed according to the size of the single sounds (Figure 1A). The sound streams consisted of instances of synthetic excerpts, representing snippets cut from exemplars of the same sound texture or a different one. Specifically, stimuli were presented in triplets: two sounds were identical (repeated; n=432), while the third one was different (novel; n=216). Compared to the repeated sounds, in one experimental context, the novel sound systematically varied for its local features, while in the other for its summary statistics (Figure 1B). That is, in the Local Features experiment, the third sound was drawn from a synthetic exemplar of the same sound texture but derived from a different white noise sample. The original white noise sample used to initialize the synthesis is expected to affect the temporal structure and the statistical values of the sounds measured at the high temporal resolution, that is in its local properties. In the other experiment (Summary Statistics), the third sound originated from the same white noise sample used to generate the two identical repeated sounds but was constrained by a different set of statistics. The summary statistics of the imposed Sound Texture affect the fine-grained spectral density measured at low temporal resolution (see below; Figure 1C). To generalize local and summary computations to a vast pool of sounds, the excerpts used in the experiments were drawn randomly among the available ones; thus, the presentation order and stimuli were never the same between participants and blocks (Berto et al., 2022).

**Figure 1.**
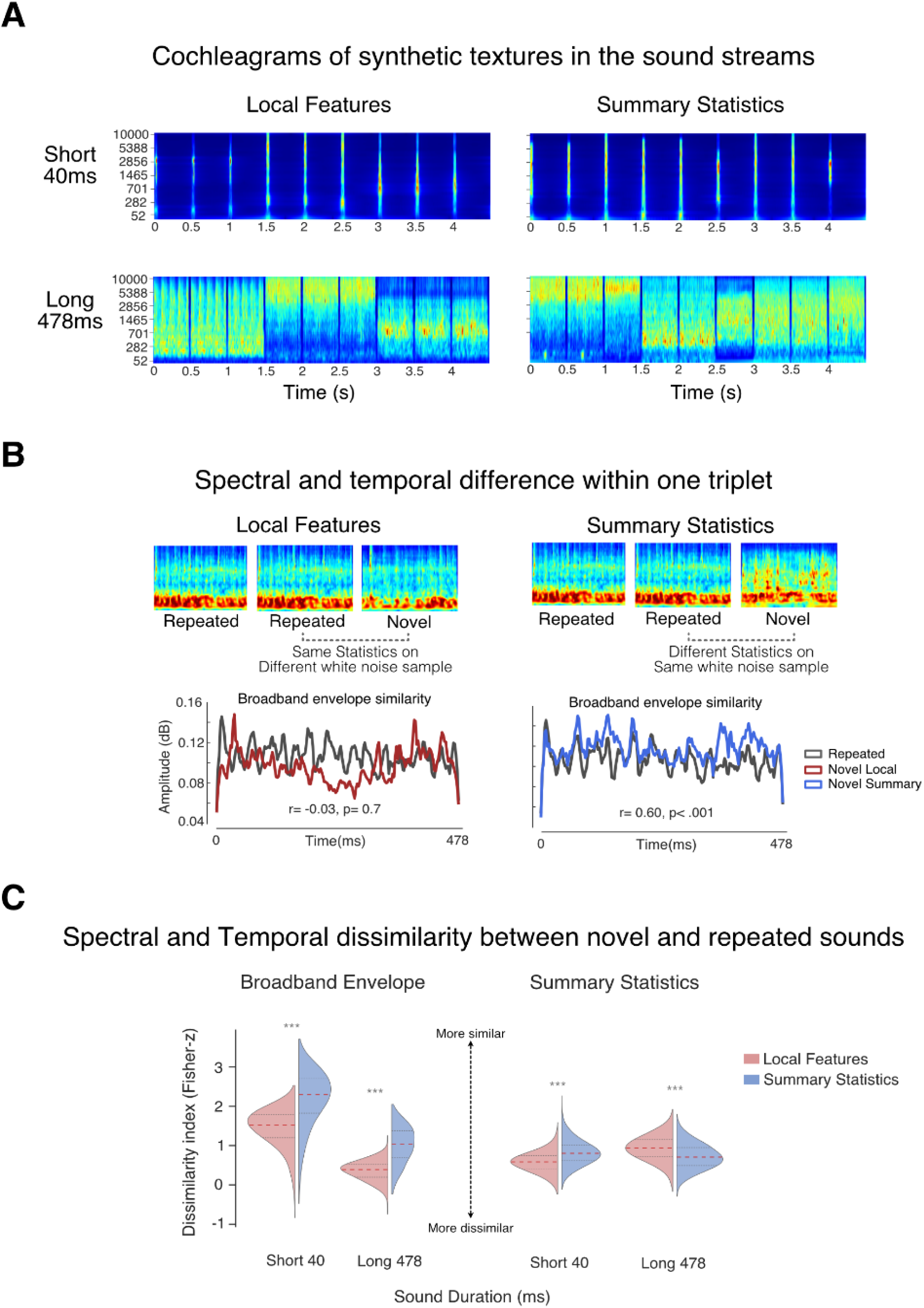
Stimuli and Experimental Protocol. (A) Cochleagrams of the first three sound pairs in a continuous stream presented to one participant. Cochleagrams were computed by filtering the sound excerpts through the Auditory Texture Model (McDermott and Simoncelli, 2011). (B) Spectral and temporal difference within a triplet. The top panel shows the cochleagrams of the sound excerpts contained in two example triplets (one for each experiment). In this example, in both experiments, the first two sounds (repeated) were a 478ms excerpt from a white noise to which we imposed the auditory statistics of the sound texture “Motorcycle idling”. In the Local features experiement, the third sound, (novel) was another exemplar of the “Motorcycle Idling” sound texture, thus it is a different white noise sample to which we impose the same statisticsin the Summary Statistics experiment, the novel sound contains the statistics from the “Idling Boat” sound texture, but it is the same white noise sample. The statistics embedded in the sounds influence the spectral density of the sound excerpts, as depicted in the cochleagram. The bottom panels display the broadband envelopes of the repeated and novel sounds in both experiments for the same three sounds displayed above (Motorcycle Idling and Idling Boat). Despite the spectral distribution between repeated and novel sounds being more similar in the Local Features experiment, the broadband envelope of novel and repeated sounds correlated more in Summary Statistics (r=0.6) than in the Local Features experiment (r=-0.03). This reveals that the temporal structure of the sounds was influenced by the original white noise sample used during synthesis, rather than the similarity of the embedded statistics, which in turn influenced the spectral content of sounds.. (C) Spectral and temporal dissimilarity between novel and repeated sound-pairs presented in the experiments.. Violin plots show the distribution of Fisher-z transformed correlation coefficients divided by experiment (Local Features and Summary Statistics) and duration (40, 478ms) for broadband envelope and auditory statistics. Within sound duration, the broadband envelopes of repeated and novel sounds were always more similar in the Summary Statistics than Local Features experiment; by contrast, auditory statistics dissimilarity showed a double dissociation according to sound duration. The violin plot outlines represent the kernel probability density (smoothed); The red dotted line is the median and gray lines are the quartiles. *** p < .001

Finally, all three sounds within a triplet had the same positional argument in the synthetic exemplar (i.e., all of them were starting and ending at the same time point along the corresponding 5s synthetic sounds they were extracted from).

Throughout the experiment, participants kept their eyes open and were instructed to look at a fixation cross appearing at the center of the screen.

The experiment was delivered by a DataPixx device (VPixx Technologies, Canada) and was programmed in MATLAB using the Psychtoolbox-3 (Brainard, 1997). Additionally, a class-based abstraction layer (https://gitlab.com/thht/o_ptb; Hartmann & Weisz, 2020) programmed on top of Psychtoolbox was used to facilitate the management of audio and triggers delivery in DataPixx.

### Spectral statistics similarities between excerpts pairs at short and long durations

To test whether the similarity in the temporal structure of repeated and novel sound excerpts was influenced by the original white noise sample used to initialize the synthesis (different in Local Features; same in Summary Statistics), we used the Auditory Texture Model (McDermott & Simoncelli, 2011; see Extended Methods in Supplementary Material) to obtain the cochleagram of each excerpt pair presented in the experiments across all participants, for each experiment (Local or Summary) and duration (40 or 478ms). Note that the same sound excerpt can appear more than once in this distribution, as we selected the precise sound sequence presented in the experiment to participants which included sounds from the same pool of available ones. Furthermore, each participant received idiosyncratic stimulation; that is, stimuli were randomly selected and paired among the available ones sharing the desired properties and sequences were never the same across participants. We then averaged over frequency channels (n=32) at each time-point to obtain the broadband envelopes. Pearson’s r*s* transformed to Fisher-z-scores were used as a metric to assess the similarity between the broadband envelopes in the sound pairs presented in the Local Features and the Summary Statistics experiment. This procedure was done for each excerpt pair presented in each experiment and duration (n=19008). The normalized coefficients in Local Features were statistically compared to the ones in Summary Statistics (two-tailed t-tests) within each sound duration. Results showed that the envelopes were always significantly more similar when the excerpts originated from the same white noise sample compared to when they did not. That is, the correlation coefficients were higher in Summary Statistics experiment compared to Local Features one, both at short (p < 0.001; Local: mean=1.45, std=0.47; Summary: mean=2.20, std=0.68) and long (p < 0.001; Local: mean=0.36, std=0.25; Summary: mean=1.02, std=0.46) durations (Figure 1C, left). On the other hand, we expected statistical values between repeated and novel sounds to be more similar depending on sound duration and experimental context. To address the statistical similarity between the couples of excerpts (repeated and novel) in the study, we selected all the sound pairs (repeated and novel) presented in the study, as done before. For each sound excerpt, we extracted the set of summary statistics using the Auditory Texture Model (McDermott & Simoncelli, 2011). We then converted the values for each statistic (envelope marginal moments and correlations, modulation power, C1, or C2) to z-scores and averaged them at each cochlear channel (n=32). The normalized averaged statistics measured from a repeated sound were correlated with the ones measured from the paired novel one. Pearson’s r*s* were transformed in Fisher-z-scores and used to assess the (dis)similarity between the statistics in the sound pairs presented in the Local Features and the Summary Statistics experiment (two-tailed t-tests within each sound duration, as before). Results showed that statistical values were significantly more dissimilar in the Local Features experiment when sounds were short (p < 0.001; Local: mean=0.58, std=0.26; Summary: mean=0.82, std=0.31) and in the Summary Statistics experiment when they were long (p < 0.001; Local: mean=0.93, std=0.32; Summary: mean=0.72, std=0.32). This dissimilarity metric (Fisher-z) is displayed in Figure 1C (right). Ultimately, these results suggest that the statistical similarity between sound excerpts depends on sound duration.

## Data analysis

### Preprocessing of MEG data

Right after the recording, a signal space separation method from the Maxfilter software (MEGIN Oy, Espoo, Finland was used to correct for different head positions across blocks within each participant and to clean channels from external interference (Taulu et al., 2005).

Preprocessing of the max-filtered MEG data was performed using MNE-Python version 1.1.1 (Gramfort et al., 2013) running on Python 3.9.16. For each participant, the signal recorded for each block was concatenated; segments contaminated, by either the beep sound or the button press, were marked for rejection. Independent component analysis (using the fast ICA method; Hyvarinen, 1999) was run to detect stereotypical artifacts (eye movements, blinks, and heartbeat). We filtered the signal below 1Hz and downsampled it to 250Hz to reduce computational time. To account for the presence of different type of channels (e.g., EOG, ECG, MEG sensors), and therefore different units (volts or tesla), the data from each channel type were pre-whitened, that is, they were scaled by the standard deviation across all channels (z-standardized). ICA was fit to the data using all 306 channels, without any prior dimensionality reduction. A specific number of components was then automatically selected for each subject based on the given variance level and accounting for rank deficiency of the data. To select which components to remove, we used a semi-automatic template-matching procedure (Campos Viola et al., 2009). Specifically, we manually selected a component (template) from one subject which best represented each artifact (one template for the eyes and one for the heart; in this case, we selected the 2 templates from 2 different subjects). Templates were chosen after visual inspection of the topography of the inverse weights and double-checked by looking at the IC activation scroll. To have additional proof that the selected templates represented eye and cardiac artifacts, we also performed Pearson correlation between all components for the selected subjects and their EOG and ECG components. Then we used the corrmap function (implemented in MNE-Python) to detect all the other components in our dataset (across all subjects) which correlated with the template above a certain threshold (0.85 for eye; 0.60 for heart). Thresholds were selected so that corrmap could find at least one IC for each subject that matches the template. Implementing the template-matching method allowed to improve and speed up bad component detection even for those subjects where EOG and ECG channels were compromised. After the components to exclude were selected, they were removed from the original unfiltered, full-resolution (1000Hz) dataset. On average, 2.68 components per subject were removed (std=0.82; range=2-5). The signal was then lowpass filtered at 40Hz (one-pass, zero-phase, non-causal FIR filter; windowed time-domain design, window type=hamming; passband ripple=0.0194; stopband attenuation=53dB; upper passband edge=40Hz; upper transition bandwidth=10Hz; -6dB cutoff frequency=45Hz; filter length=331 samples) and highpass filtered at 0.1Hz (one-pass, zero-phase, non-causal FIR filter; windowed time-domain design, window type=hamming; passband ripple=0.0194; stopband attenuation=53dB; lower passband edge=0.10Hz; lower transition bandwidth=0.10Hz; -6dB cutoff frequency=0.05Hz; filter length=33001 samples).

### Epoching

Data was epoched into segments from -100 to 500ms from the onset of the novel stimulus and downsampled to 250Hz. To correct for the physical delay of 16.5ms in sound delivery due to transmission in the air-conducting earphones, the signal was shifted in time by 0.0165 seconds. Epochs were averaged to compute event-related fields (ERFs) and baseline correction was applied by subtracting the averaged pre-stimulus period activation from -100 to 0ms from each time point post-stimulus.

The number of epochs was equalized between different experimental conditions (the difference depended on the number of beeps presented in the block, from 1 to 3).

### Source Localization

A semi-automatic co-registration pipeline was used to compute the head models. This approach has been shown to lead to comparable results as compared to manual co-registration ones (Houck & Claus, 2020). Participants’ head shapes were co-registered to a “standard” model created by the combination of 40 MRI scans of real brains (fsaverage; Fischl, 2012). For each subject, co-registration was performed in 4 steps. In step 1, we loaded the max-filtered head-shape points (HSP) and defined the fsaverage template from Freesurfer (Fischl, 2012). In step 2, we estimated three fiducial points from the fsaverage template and aligned them to the fiducials in the digitized HSP. In step 3, the outcome was refined using the Iterative Closest Point (ICP; Besl & McKay, 1992) algorithm with a small number of iterations (n=6). Any point in the HSP with a distance from the scalp above 5mm was considered an outlier and omitted. Finally, in step 4, ICP was performed once again, with a higher number of iterations (n=20) for the final co-registration fit. Head models and sensors were displayed at the end of any stage for visual inspection, to evaluate the quality of the fit and make sure that the head model was comprised within the sensors and did not fall outside. For three participants, co-registration was successful after step 3 and worsened at step 4, which was then omitted. For one participant, co-registration was effective at step 2, so steps 3 and 4 were both ignored. Finally, for one participant, co-registration failed at all steps, thus the subject was excluded from further data analysis.

The co-registered head models were used to calculate the forward solution.

First, we computed the single-layer boundary element model (BEM; (Akalin-Acar & Gençer, 2004) to create a BEM solution for the fsaverage template brain. Second, to define the position and orientation of the sources, we create a decimated dipole grid on the white matter surface using an icosahedron subdivision (ico-4) which included 2562 sources for hemispheres. Assuming a cortical surface of 1000cm^2^, the approximate spacing between the grid points was 6.2mm, and each voxel occupied 39mm^2^ of the cortical surface.

To control for field patterns related to noisy sources (e.g., human-based or environmental), we weighted each channel by a full noise covariance matrix computed on the nearest empty room recording. That is, prior to any daily data acquisition, 2 minutes of empty room measurements (data collected with no subject) were recorded in the MEG; depending on the date of testing, the closest measurement was found for each subject and preprocessed in the exact same way (max filtered; high-pass filter: 0.1Hz; low-pass filter: 40Hz). The Noise covariance matrix and the true rank were computed from the empty room measurement and used to calculate the inverse solution based on the L2 Minimum Norm Estimates.

Finally, Source Time Courses were estimated by applying the inverse operator (lamba=2; SNR=3; method=minimum norm estimates (MNE); Hämäläinen & Ilmoniemi, 1994) on the evoked data and pre-stimulus periods (from -100 to 0ms) were cropped to reduce computational space and time (as statistics were performed at latencies above 0; see Statistical Analyses below).

## Statistical Analyses

Since repeated sounds were always identical in both Local features and Summary Statistics, assessing whether the responses to the novel sounds differed between experiments would reveal if and which brain correlates discriminate based on local features and summary statistics changes. Thus, we measured statistical differences between experiments (Local vs. Summary) and within duration (Short 40ms and Long 478ms) for the evoked response to the novel sounds in source space.

Statistics were assessed using eelbrain, a Python toolkit (Brodbeck et al., 2022). Source estimates were imported in eelbrain from MNE-Python for each subject, experiment, and duration. A cluster-based permutation test for paired samples was used to test for differences in source activation at any time-point (Maris & Oostenveld, 2007). The procedure works as follows. We ran a two-tailed t-test, with a critical alpha of 0.05 at all latencies (from 0 to 500ms) and all sources. To form spatial clusters, an adjacency matrix is computed based on the fsaverage template brain and the ico-4 source space. To address the multiple comparison problem, we used a threshold-free cluster enhancement procedure (TFCE; Smith & Nichols, 2009). For every permutation of the data (n=10000), we computed a parameter map and processed it with the cluster enhancement algorithm in steps of 0.1. Subsequently, for each permutation, the maximum value of the test statistic was measured across the whole map to obtain the distribution of t-values under the null hypothesis. Based on its position in the distribution, each data point in the parameter map is assigned a p-value. To avoid considering spurious results, we assumed a stringent spatiotemporal criterion, and accepted only clusters of activation perduring at least 20ms and including a minimum of 20 sources.

## Results

In this study, we addressed whether, depending on sound duration, a change in specific sound properties (local details or summary statistics) engaged different brain correlates. That is, we tested if the auditory cortex is organized in sub-regions for selective computations based on the temporal resolution at which the sound change has occurred. To this aim, we compared the response to novel sounds between Local Features and Summary Statistics experiments within short (40ms) and long (478ms) sound duration. Any dissociation according to sound duration would indicate that the brain is endowed with distinct neural substrates for the processing of local details and summary statistics. This was clearly supported by our findings.

Multiple regions of the right hemisphere were selectively engaged for Local Feature changes occurring with short duration. Indeed, when comparing the brain response to novel sounds between Local Features and Summary Statistics experiments for short sound duration (40ms), we found a large cluster of higher activation for the Local Features experiment selectively in the right hemisphere (p < 0.001; max t-value=8.06; Cohen’s d=1.22; Figure 2A). The cluster started 148ms after stimulus onset, lasted until 224ms (duration: 76ms), and spatially comprised multiple (177) voxels of cortical sources. The cluster involved voxels embedded in several cortical regions in the auditory cortex, specifically: the superior temporal sulcus (STS) and gyrus (STG), the anterior transverse temporal gyrus (Heschl gyrus; HG), the planum temporale (PT), and the posterior segment of the lateral fissure. Other regions outside of the auditory cortex included: the central operculum, the angular and supramarginal gyri (AG and SMG), the postcentral and central sulci, the superior segment of the circular sulcus of the insula, the Jensen sulcus, the interparietal sulcus, and transverse parietal sulci (Table 1A).

**Figure 2.**
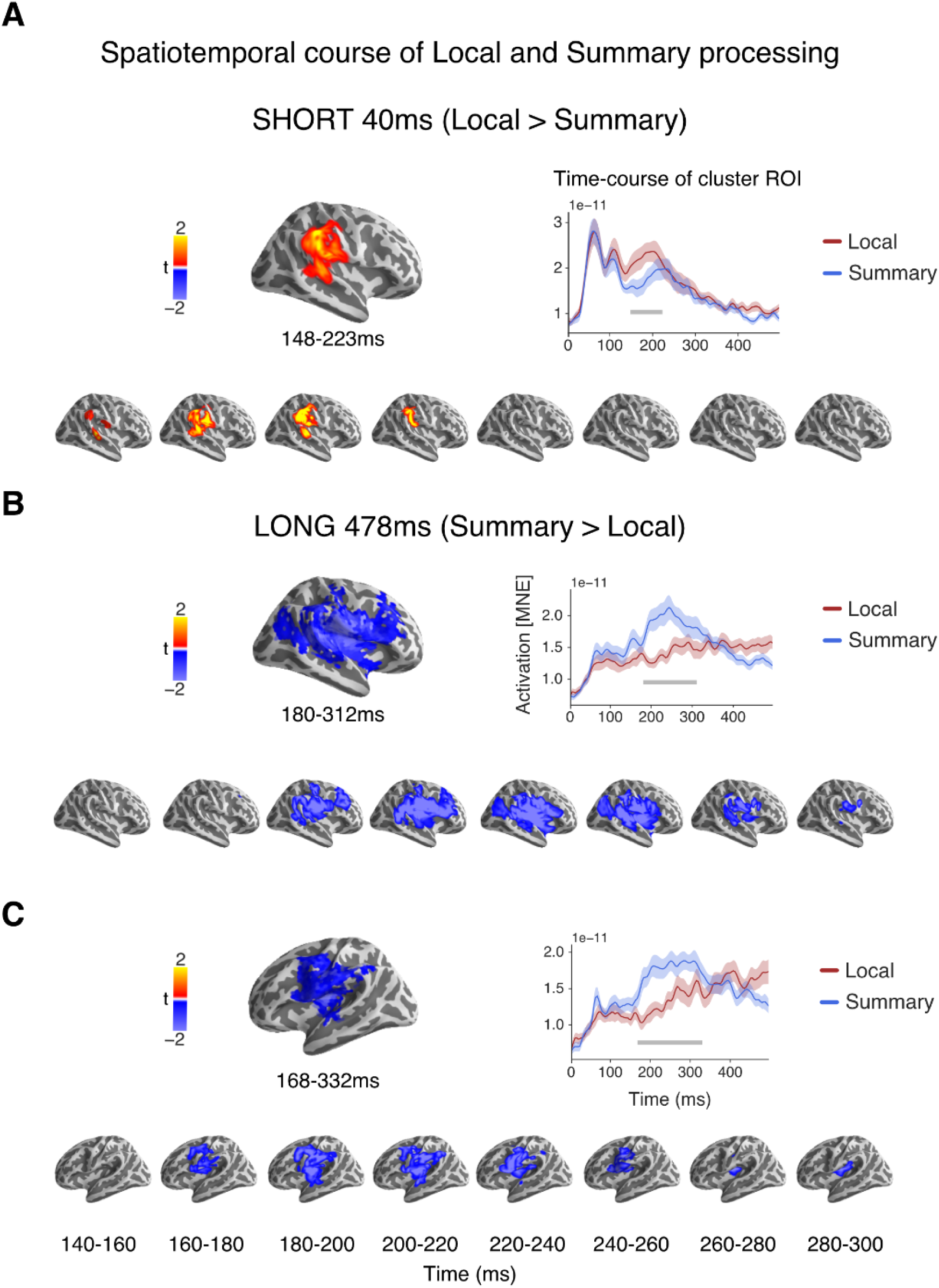
Clusters of different activation between experiments at short (40ms) and long (478ms) durations. (A) Source localization of regions whose activity was higher for Local Features as compared to Summary Statistics when sounds were short (40ms). (B) Source localization of regions in the right hemisphere whose activity was higher for Summary Statistics than Local Features when sounds were long (478ms). (C) Source localization of regions in the left hemisphere in which activity was higher for Summary Statistics than Local Features when sounds were long (478ms). For each figure (A, B, or C), the top left panel shows the t-values of significant differences between the conditions plotted on the brain surface, at the latencies displayed below. The top right panel represents the time course of activation (computed with MNE) extracted from the selected regions of interest pertaining to the cluster of activation shown on the left for both experiments (Local or Summary). Shaded areas are the standard error of the mean (SEM). Gray lines represent latencies where activations significantly differ between experiments (p < 0.05). The bottom panel displays the spatiotemporal course of cluster t-maps plotted on the brain surface at latencies displayed below.

**Table 1.**
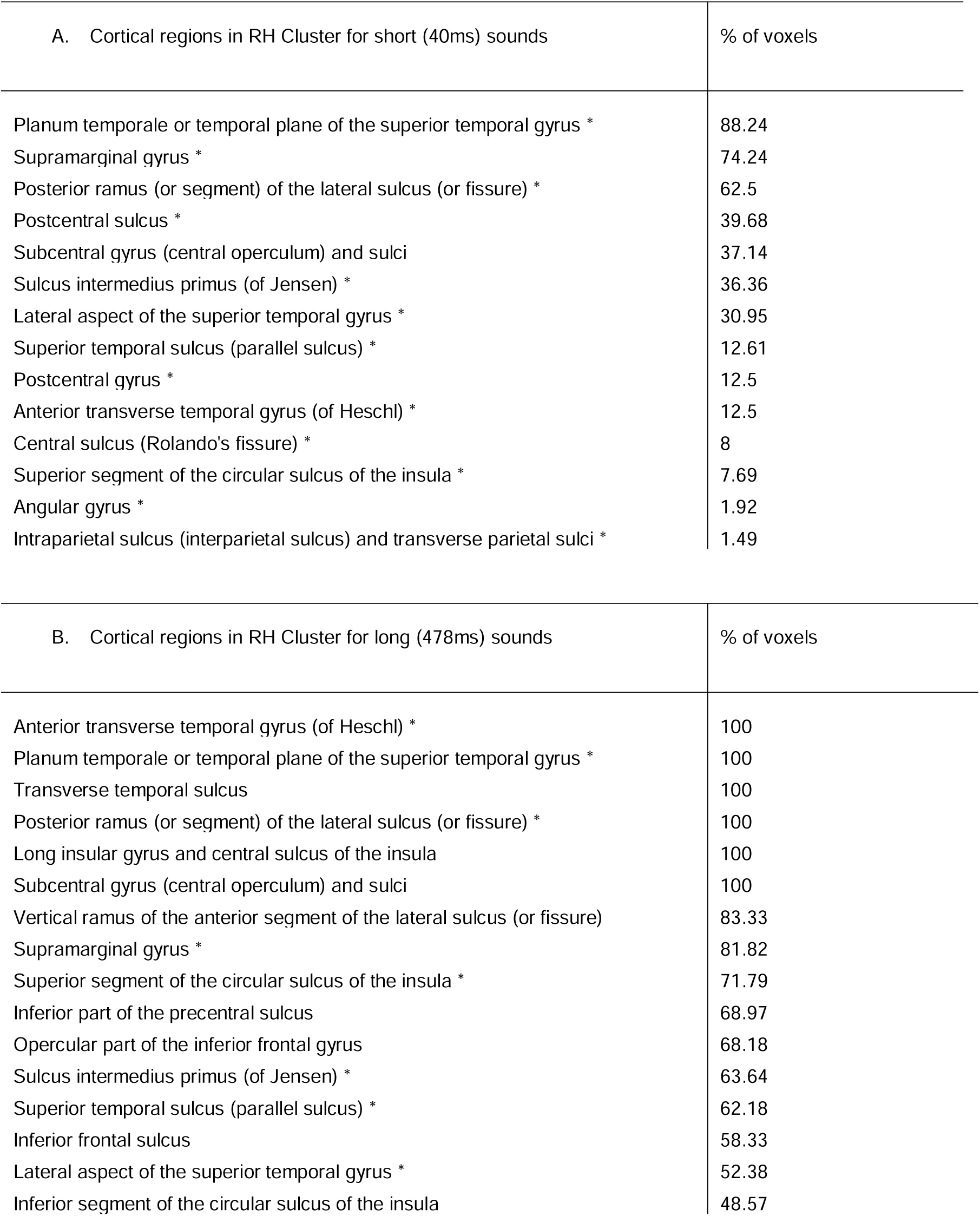
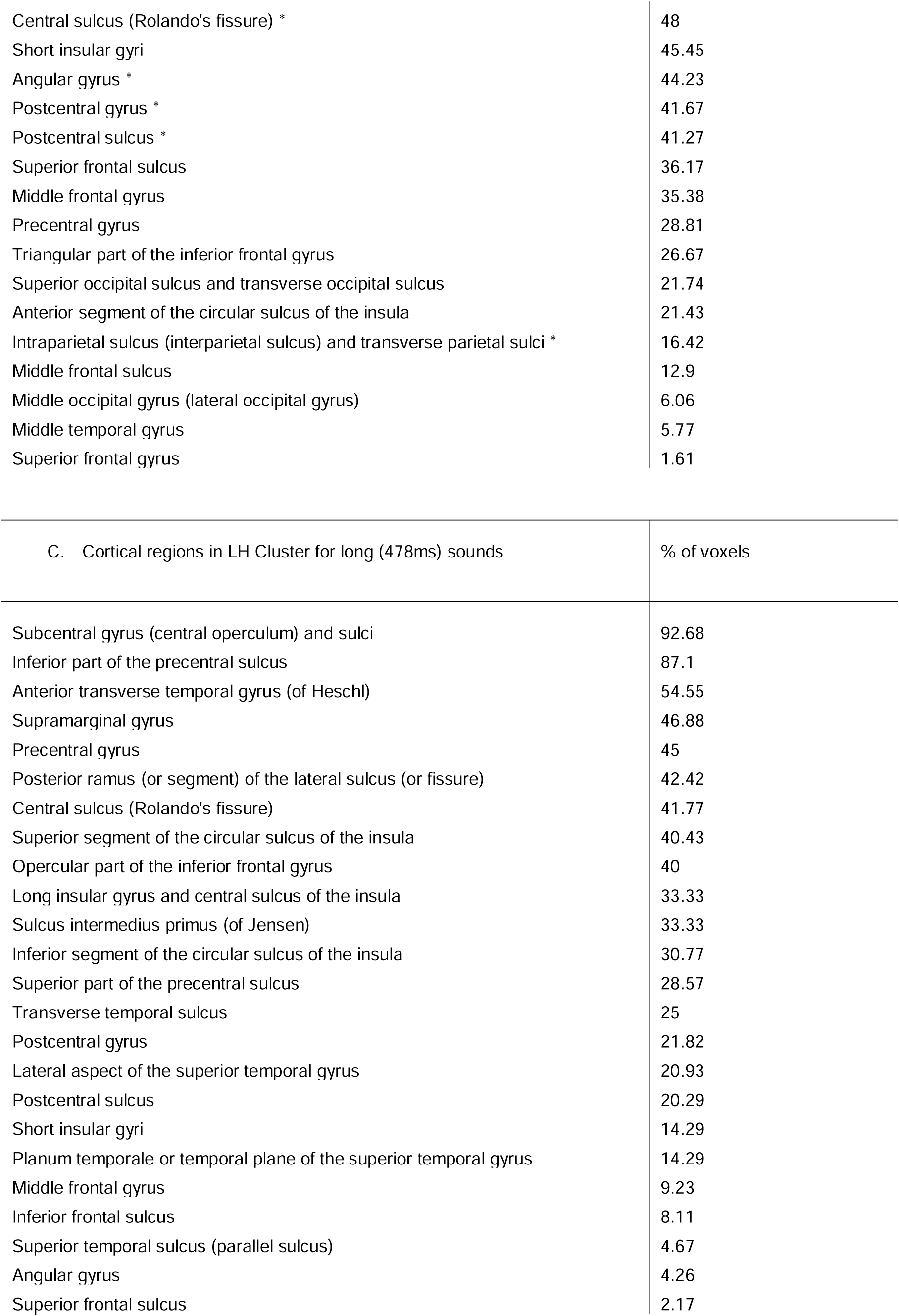
List of cortical regions in the clusters.

Conversely, changes associated with Summary Statistics occurring for long sounds (478ms) engaged both left and right hemispheres. For sounds of this duration, two large regions were significantly more responsive to novel sounds in the Summary Statistics experiment as compared to the Local Features one: (i) one in the right hemisphere and (ii) the other in the left one (Figure 2B, C).

i. The cluster found in the right hemisphere (p < 0.001; t-max=6.62; Cohen’s d=1.11) perdured from 180 to 312ms after stimulus onset (duration=132ms) and comprised 589 voxels of cortical sources. Some of the sources overlapped with the cluster observed in response to local features at the short duration, specifically around the primary auditory cortex, Heschl gyrus (HG), some temporal auditory regions such as STG, PT, STS, the lateral fissure, and other areas such as the Jensen sulcus, AG, SMG, the circular sulcus of the insula, the interparietal and transverse parietal sulci, and the central and postcentral sulci. The other non-overlapping regions comprised frontal areas, including portions of the superior, middle, and inferior frontal sulci (S-M-IFS) and gyri (S-M-IFG), and the inferior part of the precentral sulcus and gyrus; some temporal regions such as the transverse temporal sulcus, the middle temporal gyrus (MTG), the lateral fissure, and several parts of the insular cortex and central operculum; finally, some posterior regions in the occipital lobe (Table 1B).
ii. The other large cluster was found in the left hemisphere (p-value=0.004; t-max=5.77; Cohen’s d=1.13; Figure 2C). The left localized cluster started 168ms after stimulus onset and lasted until 332ms (duration=164ms). It included 289 voxels of cortical space, located mainly over the frontal lobe. In particular, the activation differed mostly over the SFS, IFS, IFG, MFG, the precentral, central, and postcentral sulci and gyri, while the cluster included fewer voxels in the temporal and parietal lobes, mainly over smaller portions of STS, STG, PT, HG, the transverse temporal sulcus, the Jensen sulcus, the insular cortex, the lateral fissure, the central operculum, and the SMG and AG (Table 1C).

The percentage of significant voxels for each region is displayed in Table 1. Noticeably, the cluster in the right hemisphere observed in response to summary statistics (long sounds) covered 100% of the cortical space in the primary auditory cortex (see Table 1A), suggesting that processing a change in summary statistics compared to one in local features requires a greater engagement of the right auditory cortex. Importantly, the areas in the right hemisphere responding to changes in local features when sounds were short and summary statistics when sounds were long, overlapped over auditory regions (Table 1A, B). That is, to some extent, the same regions in the right auditory cortex seemed to be involved in local features or summary statistics processing depending on sound duration. However, brain responses to summary statistics (long sounds) were more extended and involved anterior temporal areas and frontal regions as well.

The latencies of the two effects observed in the right hemisphere were also different, with the response to local features changes (short sounds) preceding in time (32ms before) the response to summary statistics with long sounds (Figure 1). These findings suggest that the right auditory cortex is differently engaged by one computation or the other based on sound duration; moreover, detecting a change in summary statistics, compared to local features, engages a broader network of frontotemporal regions outside auditory core ones. Conversely, the left hemisphere was involved selectively in the processing of summary statistics at long durations. The response to summary statistics predominantly involved frontotemporal areas and to a less extent primary auditory regions. When considering its temporal dynamic, the left hemisphere response to summary statistic changes (Figure 2C) started somewhere in between the responses observed for local and summary statistics changes in the right hemispheres (Figures 2A and B) and lasted longer.

Overall, these findings suggest that strong hemispheric asymmetries exist in processing Local or Summary acoustic information and that these computations rely on partially different brain sources. Specifically, when sound excerpts were short, a change in local features, with unchanged summary statistics, involved only the right hemisphere, at the level of the auditory cortex (Figure 2A). As more information was provided (e.g., stimulus duration increases), a change in summary statistics (compared to local features only) elicited higher bilateral responses in a broad network of frontotemporal areas (Figure 2B, C).

**Table 1**. In this table, we show the extended labels (from Destrieux et al., 2010) of cortical regions associated with the voxels in the three clusters with higher activation in Local Features for short (40ms) sound duration and Summary Statistics for long (478ms) sound duration. For each region, we specify the percentage of cortical surface that was included in the cluster (for instance, when the percentage of voxels is 100, the entire surface of the parcellated area is embedded in the cluster; when it is 50, only half of the voxels in that specific region belongs to the cluster, and so on). Stars * indicate the regions in the right hemisphere (RH) that are shared between the first and second clusters (A and B; See also Figure 2A, B). Cortical regions are labeled using the parcellation atlas available in the FreeSurfer package (Freesurfer v7.3.2, aparc.a2009s; Destrieux et al., 2010).

## Discussion

The aim of this study was to uncover whether the human brain is endowed with specialized neural architectures for the computation of local and summary statistics of stationary sounds.

Here, we employed a validated experimental protocol (see Berto et al., 2022) and used MEG to estimate cortical sources of these two auditory modes of functioning. We presented triplets of sounds at a constant stimulation rate, with the first two elements of the triplet being repeated and the third systematically varying for either its local temporal details or summary statistics. Since we expected local and summary processing to rely on different sound properties (i.e., impulse amplitude modulations in time and averaged spectral statistics, respectively) we hypothesized they could also rely on different auditory networks.

The predictions were confirmed by our findings. First, we found a cluster of greater activation for the Local Features experiment than the Summary Statistics one, selectively for short sounds. The opposite was found for long sounds, with clusters of higher activation in response to a change in summary statistics as compared to local ones. The clusters did not only differ in magnitude but also in their temporal and spatial distributions and hemispheric lateralization. This clear dissociation emerging according to sound duration suggests that discriminative responses to sound changes are, at least partially, guided by stimulus properties measurable at high or low temporal resolutions. Specifically, when little information is presented, the processing of local features subtends to detecting sound changes; as sound duration increases, local features become less relevant, and deviant discrimination relies on summary statistics. Interestingly, these differences in activation emerge automatically and do not require participants to be involved in a task. That is, they are engaged by a systematic and regular change in incoming stimulation. Interestingly, such responses are not stimulus-specific but represent a general mechanism of encoding the selective type of sound change. In fact, in our study, all sounds between triplets were different, the only constant was the change in local features or statistics between novel and repeated sounds. It is also worth noting that by using synthetic sounds, we ensured that the change in either temporal features or statistics was the only difference between experiments, thus any effect we found could not be explained by sound duration or other confounds.

### Functional cortical specialization for local features and summary statistics

Source estimates results confirmed the expected dissociation and highlighted partially different correlates in response to sound changes based on local features or summary statistics.

For short (40ms) sounds, a change in local features, compared to a change in summary statistics, elicited higher activations in the right hemisphere, involving several regions, including the primary auditory cortex (HG), the PT, the middle portion of the STG, and the SMG (Figure 2A; Table 2A). The higher response in these areas was sustained for several milliseconds (148-224ms) and suggested an increased sensitivity of the right auditory cortex to changes in the local details of brief sound excerpts. In line with this finding, a recent study highlighted the contribution of both HG and STG in assessing the perceived dissimilarity of sounds and suggested an active role of these regions in the processing of fine-grained acoustic details (Giordano et al., 2023). Interestingly, in the context of complex-sounds perception, the PT has been described as a computational hub devoted to segregating spectrotemporal features from complex auditory patterns (Griffiths & Warren, 2002). This includes, among others, computations relying on basic acoustic properties, such as amplitude (Giraud et al., 2000) and frequency (Hall et al., 2002) modulations. Finally, the involvement of the right SMG is in line with previous evidence appointing this brain region as part of a network involved in auditory memory (Jerde et al., 2011) and rhythmic perception (Schaal et al., 2017). Presenting the novel sounds at a regular pace creates a rhythm in the sound stream, and higher activations in the right PT and SMG may reflect the perception of a pattern influenced by the acoustic dissimilarity between repeated and novel sound excerpts.

Conversely, for long (478ms) sounds, the right hemisphere was more responsive to the processing of summary statistics changes than local ones. Such differential activation involved a broad frontotemporal network comprising several auditory regions (Figure 2B; Table 1B). Compared to the activation in favor to local features changes in short sounds, the right hemisphere activation favoring summary statistics at long durations also involved anterior and ventral temporal regions. This is in line with previous evidence showing that regions located posterior-laterally to HG encode broadband spectral information with high temporal precision, while regions located antero-ventrally preferably process fine-grained spectral information at a lower temporal resolution (Giordano et al., 2023; Kumar et al., 2007; Santoro et al., 2014). That is, as more information is accumulated, spectral content is summarized in time; in turn, the difference in activation between local and summary statistics extends to the anterior and ventral parts of the auditory cortex. The activation also spread along the STS, especially reaching its posterior part, suggesting the emergence of abstraction mechanisms (Warren et al., 2005) in response to a change in summary statistics. The STG is part of the ventral stream (or “what” pathway) for sound recognition; the posterior part, especially laterally to HG, is bilaterally activated by environmental and natural sounds (DeWitt & Rauschecker, 2012; Doehrmann et al., 2008). Moreover, different groups of neurons in STG may be responding to selective properties of sounds; such activations are then processed in higher-order areas (possibly ventrolateral prefrontal cortex; Hjortkjær et al., 2018) to resolve the category-specificity of the information (Giordano et al., 2023). Coherently, not only we observed frontal activations in the right hemisphere but also in the left one. Specifically, we found that the left hemisphere was more engaged for a change in summary statistics selectively at long sound duration. Such higher activation comprised fewer auditory regions and more areas located in the inferior frontal lobe (Figure 2C). Inferior frontal regions, known as “auditory-related areas”, have been associated with sound-object selection and are responsive to both targets and non-targets sounds, if relevant (Steinschneider et al., 2014). Previous studies suggested that local information supports sound differentiation while the emergence of summary statistics may support sound recognition (e.g., Zhai et al., 2020). In the same vein, the automatic tuning to systematic changes in generative statistics would also signify a systematic change in sound category and involve a network of activation spreading towards non-core auditory regions and frontal auditory-related areas.

Overall, these findings showed that the processing of acoustic changes at high or low temporal resolutions relies on overlapping regions in the right hemisphere but also engages different cortical areas (across the two hemispheres and extending to more anterior regions), suggesting a functional specialization in the auditory pathway for the processing of local and summary representations.

### Hemispheric specialization of auditory computations at high and low temporal resolution

Previous studies have investigated hemispheric specialization in auditory processing (i.e., Flinker et al., 2019; Zatorre & Belin, 2001). Auditory regions of one hemisphere can be more sensitive to specific sound properties than their contralateral homologue. For instance, the left hemisphere is more prone to process temporal modulations, while the right one favors the spectral content of sounds (Zatorre & Belin, 2001). Other studies showed that the left hemisphere specialized in processing information at high temporal resolutions while the right hemisphere at slower temporal rates (Flinker et al., 2019). A recent study in the mouse model demonstrated that the activation of single neurons in the right auditory cortex persists longer than their homologue in the left hemisphere (Neophytou et al., 2022). This could underlie a right hemispheric specialization in retaining brief auditory signals in memory, consistent with previous findings in humans showing longer integration windows in the right STG than in the left one (Arnal et al., 2015). Hemispheric specializations for specific auditory properties also emerged in areas outside the core auditory regions and the temporal lobe, for example, the SMG (Schaal et al., 2013, 2017).

Taken together, these findings showed that hemispheric selectivity for sound processing strictly depends on the type of stimulus that is presented, its spectral components, the administration protocol (e.g., fast or slow presentation schemes), and the task demands (i.e., rhythm or pitch perception). However, the exact computations performed by each hemisphere remain unclear, as does the general and practical meaning of such lateralization effects.

In this study, by using a systematic synthesis approach, we could directly tackle specific auditory computations (local and summary processing) while generalizing to a vast category of stimuli. That is, we presented different sound textures paired according to their intrinsic local details or summary statistics similarity, so we could measure the computations underlying discriminations based on local features and summary statistics, disregarding the characteristics of the specific sounds (i.e., pitch, acoustic frequency distribution).

Results showed that the right and left hemispheres are engaged differently depending on which computation is involved in the discrimination. When sounds are short, the processing relies on local features and involves auditory regions located in the right hemisphere, in line with evidence suggesting a right dominance for retaining in memory acoustic features (Neophytou et al., 2022).

As more information is presented, we expect a switch in computations to face acoustic overload. That is, when sound duration increases, information is averaged into summary statistics. In agreement, higher activations were expected when sounds varied for summary statistics than local features only. This was observed in both hemispheres; in the right hemisphere, similar regions observed for local features processing in short sounds were involved in computing discrimination of summary statistics in long sounds, together with non-overlapping ones. This finding implicates the right hemisphere as more involved at general stages of processing, with its computational role adapting to sound duration. In the left hemisphere, the response between experiments differed in a network of frontotemporal regions, implicating the left inferior frontal lobe selectively processes spectrotemporal information at a lower temporal rate. That is, while the right hemisphere was involved in both local and summary statistics processing, the left hemisphere was specifically engaged in the processing of summary statistics in long sounds. Interestingly, in long sounds, the activation to summary statistics in the left hemisphere precedes in time and lasts longer than the one observed in the right hemisphere. It is possible that selective auditory attention could have contributed to the left hemispheric engagement when summary statistics and sound duration could provide sufficient input to discriminate that the sound object changed. A specialization of the left hemisphere in the attentional selection of fine-grained acoustic information has previously been reported (e.g., (Bidet-Caulet et al., 2007). Altogether, these findings suggest that the right auditory cortex may be more involved in general operations subtending to both local and summary computations, such as local features retention and their subsequent integration into compact representations. On the other hand, the left hemisphere may have specialized at a higher level of processing. For instance, it might be involved in top-down regulations to guide the averaging of statistics, to evaluate if features belong to a different sound-object when a change in statistics is detected, or, in general, to select which computations to entrain (local or summary) based on monitoring the amount of entering information.

## Conclusion

Our results revealed a clear dissociation in the processing of local features and summary statistics according to sound duration. These findings allowed us to uncover the neural regions associated with these two modes of acoustic representation of stationary sounds. We highlighted how the right hemisphere developed a similar architecture to perform sound discrimination based on local features and summary statistics but possesses the ability to adapt its computations based on sound duration. On the other hand, the left hemisphere was selectively involved at a higher level of processing. By combining computational auditory modeling, a systematic approach to synthesizing sounds, and the measurement of brain activity using MEG, we provide evidence concerning the foundations of different sensory computations and their associated cortical networks.

Overall results revealed that the human brain is endowed with partially overlapping and partially specialized neural architectures for the computation of local and summary statistics of stationary sounds.

## Acknowledgments

We thank the research team in Salzburg for sharing their expertise and resources and all the master students that helped with data collection. A special mention to Manfred Seifter who assisted during all the MEG measurements.

## References

Akalin-Acar, Z., & Gençer, N. G. (2004). An advanced boundary element method (BEM) implementation for the forward problem of electromagnetic source imaging. Physics in Medicine & Biology, 49(21), 5011. https://doi.org/10.1088/0031-9155/49/21/012

Arnal, L. H., Poeppel, D., & Giraud, A. (2015). Chapter 5—Temporal coding in the auditory cortex. In M. J. Aminoff, F. Boller, & D. F. Swaab (Eds.), Handbook of Clinical Neurology (Vol. 129, pp. 85–98). Elsevier. https://doi.org/10.1016/B978-0-444-62630-1.00005-6

Berto, M., Ricciardi, E., Pietrini, P., & Bottari, D. (2021). Interactions between auditory statistics processing and visual experience emerge only in late development. IScience, 24(11), 103383. https://doi.org/10.1016/j.isci.2021.103383

Berto, M., Ricciardi, E., Pietrini, P., Weisz, N., & Bottari, D. (2022). Distinguishing fine structure and summary representation of sound textures from neural activit. https://doi.org/10.1101/2022.03.17.484757

Besl, P. J., & McKay, N. D. (1992). Method for registration of 3-D shapes. Sensor Fusion IV: Control Paradigms and Data Structures, 1611, 586–606. https://doi.org/10.1117/12.57955

Bidet-Caulet, A., Fischer, C., Besle, J., Aguera, P.-E., Giard, M.-H., & Bertrand, O. (2007). Effects of Selective Attention on the Electrophysiological Representation of Concurrent Sounds in the Human Auditory Cortex. Journal of Neuroscience, 27(35), 9252–9261. https://doi.org/10.1523/JNEUROSCI.1402-07.2007

Brainard, D. H. (1997). The Psychophysics Toolbox. Spatial Vision, 10(4), 433–436.

Brodbeck, C., Das, P., Gillis, M., Kulasingham, J. P., Bhattasali, S., Gaston, P., Resnik, P., & Simon, J. Z. (2022). Eelbrain: A Python toolkit for time-continuous analysis with temporal response functions (p. 2021.08.01.454687). bioRxiv. https://doi.org/10.1101/2021.08.01.454687

Campos Viola, F., Thorne, J., Edmonds, B., Schneider, T., Eichele, T., & Debener, S. (2009). Semi-automatic identification of independent components representing EEG artifact. Clinical Neurophysiology, 120(5), 868–877. https://doi.org/10.1016/j.clinph.2009.01.015

Destrieux, C., Fischl, B., Dale, A., & Halgren, E. (2010). Automatic parcellation of human cortical gyri and sulci using standard anatomical nomenclature. NeuroImage, 53(1), 1–15. https://doi.org/10.1016/j.neuroimage.2010.06.010

DeWitt, I., & Rauschecker, J. P. (2012). Phoneme and word recognition in the auditory ventral stream. Proceedings of the National Academy of Sciences, 109(8), E505–E514. https://doi.org/10.1073/pnas.1113427109

Doehrmann, O., Naumer, M. J., Volz, S., Kaiser, J., & Altmann, C. F. (2008). Probing category selectivity for environmental sounds in the human auditory brain. Neuropsychologia, 46(11), 2776–2786. https://doi.org/10.1016/j.neuropsychologia.2008.05.011

Fischl, B. (2012). FreeSurfer. NeuroImage, 62(2), 774–781. https://doi.org/10.1016/j.neuroimage.2012.01.021

Flinker, A., Doyle, W. K., Mehta, A. D., Devinsky, O., & Poeppel, D. (2019). Spectrotemporal modulation provides a unifying framework for auditory cortical asymmetries. Nature Human Behaviour, 3(4), Article 4. https://doi.org/10.1038/s41562-019-0548-z

Giordano, B. L., Esposito, M., Valente, G., & Formisano, E. (2023). Intermediate acoustic-to-semantic representations link behavioral and neural responses to natural sounds. Nature Neuroscience, 26(4), Article 4. https://doi.org/10.1038/s41593-023-01285-9

Giraud, A.-L., Lorenzi, C., Ashburner, J., Wable, J., Johnsrude, I., Frackowiak, R., & Kleinschmidt, A. (2000). Representation of the Temporal Envelope of Sounds in the Human Brain. Journal of Neurophysiology, 84(3), 1588–1598. https://doi.org/10.1152/jn.2000.84.3.1588

Gramfort, A., Luessi, M., Larson, E., Engemann, D., Strohmeier, D., Brodbeck, C., Goj, R., Jas, M., Brooks, T., Parkkonen, L., & Hämäläinen, M. (2013). MEG and EEG data analysis with MNE-Python. Frontiers in Neuroscience, 7. https://www.frontiersin.org/articles/10.3389/fnins.2013.00267

Griffiths, T. D., & Warren, J. D. (2002). The planum temporale as a computational hub. Trends in Neurosciences, 25(7), 348–353. https://doi.org/10.1016/S0166-2236(02)02191-4

Hall, D. A., Johnsrude, I. S., Haggard, M. P., Palmer, A. R., Akeroyd, M. A., & Summerfield, A. Q. (2002). Spectral and Temporal Processing in Human Auditory Cortex. Cerebral Cortex, 12(2), 140–149. https://doi.org/10.1093/cercor/12.2.140

Hämäläinen, M. S., & Ilmoniemi, R. J. (1994). Interpreting magnetic fields of the brain: Minimum norm estimates. Medical & Biological Engineering & Computing, 32(1), 35–42. https://doi.org/10.1007/BF02512476

Hartmann, T., & Weisz, N. (2020). An Introduction to the Objective Psychophysics Toolbox. Frontiers in Psychology, 11. https://www.frontiersin.org/articles/10.3389/fpsyg.2020.585437

Hjortkjær, J., Kassuba, T., Madsen, K. H., Skov, M., & Siebner, H. R. (2018). Task-Modulated Cortical Representations of Natural Sound Source Categories. Cerebral Cortex, 28(1), 295–306. https://doi.org/10.1093/cercor/bhx263

Houck, J. M., & Claus, E. D. (2020). A comparison of automated and manual co-registration for magnetoencephalography. PLOS ONE, 15(4), e0232100. https://doi.org/10.1371/journal.pone.0232100

Hyvarinen, A. (1999). Fast and robust fixed-point algorithms for independent component analysis. IEEE Transactions on Neural Networks, 10(3), 626–634. https://doi.org/10.1109/72.761722

Jerde, T. A., Childs, S. K., Handy, S. T., Nagode, J. C., & Pardo, J. V. (2011). Dissociable systems of working memory for rhythm and melody. NeuroImage, 57(4), 1572–1579. https://doi.org/10.1016/j.neuroimage.2011.05.061

Maris, E., & Oostenveld, R. (2007). Nonparametric statistical testing of EEG-and MEG-data. Journal of Neuroscience Methods, 164(1), 177–190. https://doi.org/10.1016/j.jneumeth.2007.03.024

McDermott, J. H., Schemitsch, M., & Simoncelli, E. P. (2013). Summary statistics in auditory perception. Nature Neuroscience, 16(4), Article 4. https://doi.org/10.1038/nn.3347

McDermott, J. H., & Simoncelli, E. P. (2011). Sound Texture Perception via Statistics of the Auditory Periphery: Evidence from Sound Synthesis. Neuron, 71(5), 926–940. https://doi.org/10.1016/j.neuron.2011.06.032

McWalter, R., & McDermott, J. H. (2019). Illusory sound texture reveals multi-second statistical completion in auditory scene analysis. Nature Communications, 10(1), Article 1. https://doi.org/10.1038/s41467-019-12893-0

Neophytou, D., Arribas, D. M., Arora, T., Levy, R. B., Park, I. M., & Oviedo, H. V. (2022). Differences in temporal processing speeds between the right and left auditory cortex reflect the strength of recurrent synaptic connectivity. PLOS Biology, 20(10), e3001803. https://doi.org/10.1371/journal.pbio.3001803

Norman-Haignere, S. V., & McDermott, J. H. (2018). Neural responses to natural and model-matched stimuli reveal distinct computations in primary and nonprimary auditory cortex. PLOS Biology, 16(12), e2005127. https://doi.org/10.1371/journal.pbio.2005127

Santoro, R., Moerel, M., Martino, F. D., Goebel, R., Ugurbil, K., Yacoub, E., & Formisano, E. (2014). Encoding of Natural Sounds at Multiple Spectral and Temporal Resolutions in the Human Auditory Cortex. PLOS Computational Biology, 10(1), e1003412. https://doi.org/10.1371/journal.pcbi.1003412

Schaal, N. K., Pollok, B., & Banissy, M. J. (2017). Hemispheric differences between left and right supramarginal gyrus for pitch and rhythm memory. Scientific Reports, 7(1), Article 1. https://doi.org/10.1038/srep42456

Schaal, N. K., Williamson, V. J., & Banissy, M. J. (2013). Anodal transcranial direct current stimulation over the supramarginal gyrus facilitates pitch memory. European Journal of Neuroscience, 38(10), 3513–3518. https://doi.org/10.1111/ejn.12344

Smith, S. M., & Nichols, T. E. (2009). Threshold-free cluster enhancement: Addressing problems of smoothing, threshold dependence and localisation in cluster inference. NeuroImage, 44(1), 83–98. https://doi.org/10.1016/j.neuroimage.2008.03.061

Steinschneider, M., Nourski, K. V., Rhone, A. E., Kawasaki, H., Oya, H., & Howard, M. A. (2014). Differential activation of human core, non-core and auditory-related cortex during speech categorization tasks as revealed by intracranial recordings. Frontiers in Neuroscience, 8, 240. https://doi.org/10.3389/fnins.2014.00240

Taulu, S., Simola, J., & Kajola, M. (2005). Applications of the signal space separation method. IEEE Transactions on Signal Processing, 53(9), 3359–3372. https://doi.org/10.1109/TSP.2005.853302

Warren, J. D., Jennings, A. R., & Griffiths, T. D. (2005). Analysis of the spectral envelope of sounds by the human brain. NeuroImage, 24(4), 1052–1057. https://doi.org/10.1016/j.neuroimage.2004.10.031

Zatorre, R. J., & Belin, P. (2001). Spectral and Temporal Processing in Human Auditory Cortex. Cerebral Cortex, 11(10), 946–953. https://doi.org/10.1093/cercor/11.10.946

Zhai, X., Khatami, F., Sadeghi, M., He, F., Read, H. L., Stevenson, I. H., & Escabí, M. A. (2020). Distinct neural ensemble response statistics are associated with recognition and discrimination of natural sound textures. Proceedings of the National Academy of Sciences, 117(49), 31482–31493. https://doi.org/10.1073/pnas.2005644117

Zuk, N. J., Teoh, E. S., & Lalor, E. C. (2020). EEG-based classification of natural sounds reveals specialized responses to speech and music. NeuroImage, 210, 116558. https://doi.org/10.1016/j.neuroimage.2020.116558

