## Supplementary Material for "Hemispheric asymmetries in auditory cortex reflect discriminative responses to temporal details or summary statistics of stationary sounds"

This document contains:

Supplementary Text: Extended Methods
Supplementary Figure S1
Supplementary Table S1
Supplementary References

### Extended Methods

#### Auditory Texture Model

The time-averaged summary statistics were measured using the auditory texture model as in McDermott and Simoncelli (2011), whose code is available here: http://mcdermottlab.mit.edu/downloads.html. For clarity, we provide a summary of the model in the following paragraph. For a detailed description, see the original paper (McDermott & Simoncelli, 2011).

The model employed a filter bank cascade to process an input sound waveform, $x(t)$, representing an original recording of a sound texture, and extracted summary statistics from their outputs. The procedure can be divided into two processing stages.

First, to replicate the frequency analysis occurring in the cochlea, the input sounds were filtered into subbands using a bank of bandpass filters with varying center frequencies and bandwidths. The model employed 4th-order gammatone filters consisting of 32 zero-phase bandpass filters with center frequencies equally spaced on an equivalent rectangular bandwidth (ERB; Glasberg & Moore, 1990) scale between 20 and 10,000Hz. The filters had a bandwidth of 3db, comparable to the one in the human ear (McDermott & Simoncelli, 2011). The outcome of the filtering stage were cochlear subbands, the analytic signal (or fine structure) at each center frequency. The envelope of each subband was then computed via Hilbert transform. To emulate the non-linearity of the basilar membrane compression, the envelopes were elevated by a power of 0.3 (Ruggero, 1992). For computational efficiency, the subband envelopes were then downsampled to 400Hz. Overall, this represented the first processing stage resulting in the cochleagrams of sounds (displayed in Figure 1A ,B).

In the second processing stage, each cochlear envelope was convolved with a second band of filters to obtain amplitude modulation rate subbands. These modulation filters consisted of 20 half-octave spaced bandpass filters (from 0.5 to 200Hz) with a constant quality factor ($Q$) of 2 (for 3dB bandwidths). This reflects the selectivity of the human auditory system, likely a result of thalamic processing (Dau et al., 1997).

Auditory texture statistics were extracted from the cochlear envelope subbands $x_k(t)$ and the modulation subbands, $b_{k,n}(t)$, where $k$ and $n$ indexed the cochlear and modulation channels, respectively.

The computed envelope statistics included three marginal moments: (i) the mean, (ii) the coefficient of variance, (iii) and the skewness. Respectively:

$$\mu_k = \sum_t w(t)x(t) \qquad \text{(i)}$$

$$\frac{\sigma^2}{\mu_k^2} = \frac{\sum_t w(t)(x_k(t) - \mu_k)^2}{\mu_k^2} \qquad \text{(ii)}$$

$$\eta_k = \frac{\sum_t w(t)(x_k(t) - \mu_k)^3}{\sigma_k^3} \tag{iii}$$

where $\mu_k$ is the mean, $\sigma_k^2$ is the variance, $\eta_k$ is the skewness at each cochlear channel, and w(t) is a windowing function, with the constraint that $\sum_t w(t) = 1$ (McDermott & Simoncelli, 2011). The variance was normalized by the squared mean to make it dimensionless, as the skewness (McDermott & Simoncelli, 2011). These marginal moments captured the sparsity of the time-averaged subband envelopes. Note that, following the original authors recommendations, compared to the previous version of the model (McDermott & Simoncelli, 2011), our implementation omitted the Kurtosis from the marginal moments, because proven to be not very informative.

The model included pairwise correlations between each cochlear channel and the eight nearest neighbors. Cross-band correlations capture broadband events that would activate more cochlear bands at the same time (Nelken et al., 1999).
The cross-band correlation coefficient can be computed as follow:

$$c_{jk} = \frac{\sum_t w(t)(x_j(t) - \mu_k)}{\sigma_j \sigma_k}, j, k \in [1...32]$$

such that $(k - j) \in [1,2,3,5,8,11,16,21]$.

Note that to capture the envelope power at different modulation rates, the modulation subband variance was normalized by the corresponding total cochlear envelope variance.

Another fundamental measure was the power at each modulation rate, namely the modulation power. The modulation power represents the major statistics of interest concerning the modulation bands and may reflect modulation-tuned properties in the thalamic neurons response (Dau et al., 1997; Miller et al., 2002). To calculate it, the model measured the variance of each modulation subband and normalized it by the total envelope variance as follow:

$$\sigma_{k,n} = \frac{\sum_t w(t)(b_{k,n}(t) - \mu_{k,n})^2}{\sigma_k^2}, k \in [1 \dots 32], n \in [1 \dots 20]$$

Finally, the model employed octave-spaced modulation filters (McDermott & Simoncelli, 2011) to measure correlations (C1 and C2) between modulation subbands of different cochlear channels. The frequency responses of the filters (seven filters, with center frequencies in octave steps from 1.56 to 100Hz) were half-cosines on a log-scale and more broadly tuned with $Q = \sqrt{2}$ .

C1 was computed as follow:

$$C1_{jk,n} = \frac{\sum_t w(t)\tilde{b}_{j,n}(t)\tilde{b}_{k,n}(t)}{\sigma_{j,n}\sigma_{k,n}}, j \in [1...32], (k - j) \in [1,2], n \in [2...7],$$

and

$$\sigma_{j,n} = \sqrt{\sum_t w(t)\tilde{b}_{j,n}(t)^2}$$

where $\tilde{b}_{k,n}(t)$ are the resulting bands of the correlation filters.

C2 is calculated as follow:

$$C2_{k,mn} = \frac{\sum_t w(t) d^*_{k,m}(t)\alpha_{k,n}(t)}{\sigma_{k,m}\sigma_{k,n}}, k \in [1 \dots 32], m \in [1 \dots 6], (n-m) = 1$$

where * denote the complex conjugate, $\alpha$ is the analytic signal of the modulation bands comprising the response of the filter and its quadratic twin, and $d$ is the frequency doubling resulting from squaring the analytic signal (complex-valued). $\alpha$ and d are calculated as follow:

$$\alpha_{k,n}(t) = \tilde{b}_{k,n}(t) + iH\left(\tilde{b}_{k,n}(t)\right)$$

and

$$d_{k,n} = \frac{\alpha^2_{k,n}(t)}{\left\| \alpha_{k,n}(t) \right\|}$$

where $i = \sqrt{-1}$ and H is the Hilbert transform.

For each sound texture, a set of time-averaged texture statistics were measured and included in a parameter vector, $\zeta$, which was subsequently utilized to generate the synthetic sounds.

**Sound texture synthesis.**

We synthesized sound textures following the procedure described in McDermott and Simoncelli (2011). The parameter vector ($\zeta$) of time-averaged statistics was measured for each of the 7s original sound textures (n=54; listed in table S1). The sound synthesis toolbox is available online:
http://mcdermottlab.mit.edu/downloads.html.
The measured statistics were then imposed on a white noise sample (seed). The algorithm measured the statistics from the seed and iteratively adjusted them to match the ones extracted from the original sound textures. To perform this computation, we used a conjugate gradient descent based on the “minimize” function by Carl Rasmussen (embedded in the toolbox, see above; McDermott & Simoncelli, 2011) which compares the total squared error of statistics measured from the synthetic sound with those from the original signal. Specifically, the procedure monitored the convergence between the statistics of the two signals by calculating the signal-to-noise ratio (SNR). The SNR was determined by comparing the squared

error of a statistic class (summed across all statistics within that class) to the sum of the squared statistic values in that class. The iteration procedure was stopped when all statistic classes had an SNR of 30dB or greater or after reaching 60 iterations. Convergence was achieved when the average SNR of all statistic classes reached 20dB or higher (McDermott & Simoncelli, 2011).

For each sound texture, the synthesis algorithm was initialized with four different 5s white noise samples resulting in four different synthetic exemplars of the same original texture.

By seeding the synthesis algorithm with different Gaussian noises, we could generate distinct exemplars of the same sound texture, including auditory statistics which converged with sound duration but retained a different fine structure (local features).

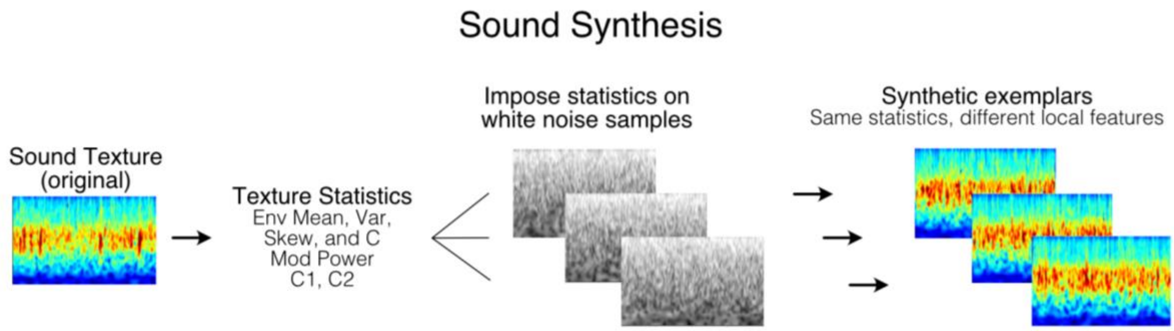


**Figure S1. Schematic representation of sound synthesis. Related to Figure 1.** An original recording of a sound texture (in this case, “Applause of a large crowd”) is filtered through the auditory texture model to extract the set of auditory texture statistics. Texture statistics are imposed on different random noise samples. The results are synthetic exemplars of the same sound textures. These sounds are constrained only by the imposed summary statistics; thus, they are perceived as the same sound object. However, each of them retains the local temporal structure of the white noise on which we initialized the synthesis. That is, they all have the same summary statistics but different local features.

### Table S1. List of the sound textures employed in the experiments.

| **Column 1:** Repeated and Novel (in Local Features) Repeated (in Summary Statistics) | **Column 2:** Only in Summary Statistics Novel |
|---|---|
| Applause large crowd | Applause big room |
| Applause 2 | Applause 1 |
| Bathroom sink | Bath being drawn |
| Bulldozer | Waterfall |
| Castanets 1 | Castanets 2 |
| Electric adding machine | Teletype city room |
| Fast running river | River running over shallows |
| Fire burning room 2 | Fire 3 |
| Fire forest inferno | Fire 3 |
| Frogs 4 | Frogs 3 |
| Frying bacon | Crunching cellophane |
| Heavy rain falling and dripping | Heavy rain on hard surface |
| Heavy rain on hard surface | Rain in woods2 |
| Horse trotting on cobblestones | Horse and buggy |
| Industrial machinery | Construction site ambience |
| Jungle rain | Rain in woods1 |
| Linotype | Teletype city room |
| Motorcycle idling | Idling boat |
| Pneumatic drills | Construction site ambience |
| Printing press | Construction site ambience |
| Radio static 2 | Radio static1 |
| Rain in woods 1 | Frogs 1 |
| Rain in woods 2 | Jungle rain |
| Rain | Rain in woods 1 |
| Rhythmic applause | Applause big room |
| River running over shallows | Applause big room |
| Shaking coins | Pouring coins1 |
| Ship anchor being raised | Pneumatic drills |
| Sparrows large, excited group | Birds in tropical forest |
| Stream near small waterfall | River running over shallows |
| Teletype city room | Teletype |
| Typewriter IBM electric | Typewriter manual |
| Water running into sink | Bathroom sink |
| Waterfall | Air conditioner |
| Frogs 3 | Frogs 1 |
| Metal lathe | Blender |
| Applause large crowd | Applause big room |

**Table S1. Related to Figure 1.** Sound textures were the same as in (Berto et al., 2021, 2022). In Local Features Discrimination, for each sound texture in column 1, two synthetic exemplars of the sound texture were selected. One was presented twice (repeated) and the other was presented as the third element of the triplet (novel). In Summary Statistics Discrimination, sound textures were paired according to perceived similarity (McDermott, Schemitsch, and Simoncelli, 2013). For each sound texture in column 1, one synthetic exemplar was selected and presented twice. Then, an exemplar of the texture from the corresponding row in column 2 was selected and used as the third element of the triplet (novel).

### Supplementary References

Berto, M., Ricciardi, E., Pietrini, P., & Bottari, D. (2021). Interactions between auditory statistics processing and visual experience emerge only in late development. IScience, 24(11), 103383. https://doi.org/10.1016/j.isci.2021.103383

Berto, M., Ricciardi, E., Pietrini, P., Weisz, N., & Bottari, D. (2022). Distinguishing fine structure and summary representation of sound textures from neural activit. https://doi.org/10.1101/2022.03.17.484757

Dau, T., Kollmeier, B., & Kohlrausch, A. (1997). Modeling auditory processing of amplitude modulation. II. Spectral and temporal integration. The Journal of the Acoustical Society of America, 102(5), 2906–2919. https://doi.org/10.1121/1.420345

Glasberg, B. R., & Moore, B. C. J. (1990). Derivation of auditory filter shapes from notched-noise data. Hearing Research, 47(1), 103–138. https://doi.org/10.1016/0378-5955(90)90170-T

McDermott, J. H., & Simoncelli, E. P. (2011). Sound Texture Perception via Statistics of the Auditory Periphery: Evidence from Sound Synthesis. Neuron, 71(5), 926–940. https://doi.org/10.1016/j.neuron.2011.06.032

Miller, L. M., Escabí, M. A., Read, H. L., & Schreiner, C. E. (2002). Spectrotemporal Receptive Fields in the Lemniscal Auditory Thalamus and Cortex. Journal of Neurophysiology, 87(1), 516–527. https://doi.org/10.1152/jn.00395.2001

Ruggero, M. A. (1992). Responses to sound of the basilar membrane of the mammalian cochlea. Current Opinion in Neurobiology, 2(4), 449–456. https://doi.org/10.1016/0959-4388(92)90179-O
